# Purification and characterization of fat body lipase from the Greater Wax Moth *Galleria mellonella* (Lepidoptera: Pyralidae)

**DOI:** 10.1101/625129

**Authors:** Rahma R.Z. Mahdy, Shaimaa A. Mo’men, Marah M. Abd El-Bar, Emad M.S. Barakat

**Author notes:** **Rahma R.Z. Mahdy:** Demonstrator at Department of Entomology, Faculty of Science, Ain Shams University, Cairo, Egypt., **Shaimaa A. Mo’men:** A Lecturer at Department of Entomology, Faculty of Science, Ain Shams University, Cairo, Egypt., **Marah M. Abd El-Bar: (The corresponding author)** Associate Professor at Department of Entomology, Faculty of Science, Ain Shams University, Cairo, Egypt (Basic), **Emad M.S. Barakat:** Professor of Insect Physiology at Department of Entomology, Faculty of Science, Ain Shams University, Cairo, Egypt.

## Abstract

Lipid mobilization and transport in insects is under investigation, especially lipases and lipophorin because of their roles in energy production and transport of lipids at flying activity. The present study has been conducted to purify intracellular fat body lipase for the first time, from last larval instar of *Galleria mellonella*. Purification methods by combination of ammonium sulfate precipitation and gel filtration using Sephadex G-100 demonstrated that the amount of protein and the specific activity of fat body lipase were 0.008633±0.000551 mg/ml and 1.5754±0.1042 μmol/min/mg protein, respectively, with a 98.9 fold purity and recovery of 50.81%. Hence, the sephadex G-100 step was more effective in purification process. SDS-PAGE and zymogram revealed that fat body lipase showed two monomers with molecular weights of 178.8 and 62.6 kDa. Furthermore biochemical characterization of fat body lipase was carried out through testing its activities against several factors such as; different temperatures, pH ranges, metal ions and inhibitors ending by determination of their kinetic parameters with the use of *p*-Nitrophenyl butyrate (PNPB) as a substrate. The highest activities of enzyme were determined at the temperature ranges of 35-37°C and 37-40°C and pH ranges of 7-9 and 7–10. The partially purified enzyme showed significant stimulation by Ca^2+^, K^+^ and Na^+^ metal ions indicating that fat body lipase is metalloproteinase. Additionally, lipase activity was strongly inhibited by some inhibitors; phenylmethylsulfony fluoride (PMSF), ethylene-diaminetetractic acid (EDTA) and ethylene glycoltetraacetic acid (EGTA) providing an evidence of presence of serine residue and activation of enzymes by metal ions. Kinetic parameters were 301.95mM K_m_ and 0.316 Umg^−1^ V_max_. By considering the purification of fat body lipase from larvae and using some inhibitors especially ion chelating agents, it is suggested to develop this study by using lipase inhibitors to reach a successful control of *Galleria mellonella* in the near future.

## Introduction

Lipids are utilized efficiently by insects as substrate for reproduction embryogenesis, metamorphosis and flight. The importance of lipids to insects had been recognized by several authors [1,2].

Lipids are large and heterogeneous group of substances which are insoluble in water but soluble in nonpolar solvents. Some lipids contain fatty acids (e.g., fats and waxes) while others lack them (e.g., terpenes). Lipids containing fatty acids involve several groups; sphinogolipids, glycerolipids, glycerophospholipids, sterol lipids, saccahrolipids, prenol lipids and polykiteds [3]. Others that contain metabolites play an important role in biological reactions [4]. Lipids containing fatty acids present as storage lipids and membrane lipids. Glycolipids and phospholipids are included in membrane lipids, while the storage lipids represented by fats in adipose tissue of animals, fat bodies in insects, or oils in seeds [5]. The principal storage site for insect lipids is the fat body, and the lipid composition of the whole insect probably reflects the lipid composition of the fat body [6,7]. Lipid content and lipid composition of the fat body are the result of various processes, including storage of dietary lipids, *de novo* synthesis, degradation and modification of fat body lipid and subsequent release for transport to sites of utilization [8].

Lipases are the enzymes responsible for the hydrolysis of lipid (triacylglycerol acylhydrolase, EC 3.1.1.3) which catalyze the hydrolysis of fatty acids ester bonds. In insects, these enzymes have key roles in utilizing, storing and transmitting lipids and also they are important in basic physiological processes of reproduction, development, defending against pathogens and oxidative stress, and pheromone signaling [2]. Insect lipases are divided into triacylglycerol lipases (TAG-lipases), alkaline and acid phosphatases in addition to phospholipases [9].

Lipases have a dynamic physiological role; the catabolism of triacylglycerols (TAGs) stored as fat depots and those from dietary lipids. Henceforth, two main groups of lipase are documented, lysosomal (intracellular) and digestive lipases [10]. Intracellular lipases are responsible of TAGs hydrolysis stored as lipid droplet, the chief endogenous source of energy [11], while digestive lipases hydrolyze TAGs in food.

The greater wax moth, *Galleria mellonella (G*. *mellonella)*, is one of the most destructive pests of honey bee colonies worldwide [12]. It causes considerable economic losses to beekeepers by damaging wax combs. The destruction of the comb will leak or contaminate stored honey and may kill bee larvae or be the cause of the spreading of honey bee diseases. However, an effective method of controlling this pest has not been developed. Physical and chemical methods are imperfect. Therefore, many studies have been conducted to find ways of controlling it. It is mandatory to study the enzymes of the insect pests in order to develop biotechnological processes to provide perfect and effective control measures [12].

Recently, lipid mobilization and transport in insects is under investigation, especially lipases and lipophorin because of their roles in energy production and transport of lipids at flying activity. Most of researches performed on insect lipase focused on midgut lipase, while little researches have been carried out on fat bodies [13,14]. Lipases have been purified from some insects, such as *Manduca sexta* [6]; *Gryllus campestris* [14]; *Locusta migratoria* [15]; and *Naranga aenescens* [16].

Detailed knowledge on the enzymatic environment of *Galleria* will provide new opportunities for a sustainable pest management to control this pest. The wax moth is a highly specialized insect, its larva feeds on beeswax and probably has a unique system for lipid transport and utilization [17].

To our knowledge, no intracellular lipase has been purified and characterized in wax moth. In our preliminary study [18] we have carried out purification and characterization of midgut lipase from *G*. *mellonella* larva. Therefore, the aim of the current study is to purify and characterize fat body lipase of *G*. *mellonella* larvae.

## Materials and Methods

### Rearing and maintenance of experimental insect

The greater wax moth, *G*. *mellonella* (L.) was obtained from Plant Protection Research Institute, Agricultural Research Center, Dokki, Cairo, Egypt. A stock colony of *G*. *mellonella* was maintained for several generations in the insectary of Entomology Department, Faculty of Science, Ain Shams University and reared on artificial diet according to [19]. Last larval stages were collected and the silk cocoon around them was removed before further experiments.

### Processing of larval tissue homogenates

Enzyme extracts of fat body tissues of *G*. *mellonella* larvae were prepared according to [20].

#### Fat body collection

The larval body was squeezed and the haemolymph was discarded. The fat body was dissected out. Pooled tissues from 50 individuals were immediately transferred to buffered saline (pH 6.9), rinsed, and weighed prior to any analysis or incubation.

#### Preparation of larval tissue homogenates

Fatbody tissues were rinsed in ice-cold buffer, placed in a pre-cooled glass homogenizer and ground in 1 ml of buffer solution (pH 7). The homogenates were separately transferred to 2 ml centrifuge tubes and centrifuged at 13,000 rpm (Human Centrifuge, TGL-16XYJ-2, 16000 rpm, Korea) for 20 min at 4°C. The supernatants were pooled and stored at −20°C for subsequent analyses.

#### Protein concentration

The protein concentration of was determined by the method of Bradford [21] by using bovine serum albumin (BSA) as a standard protein.

### Determination of lipase activity

lipase assay was carried out by using continuous spectrophotometric rate determination method as described by Tsujita et al. [22] with some modifications. 30 μl of crude extracts (from fat body) and 100 μl of p-nitrophenyl butyrate (pNPB, 50 mM), as substrate were mixed thoroughly and incubated at 37°C. For negative control tubes, samples were placed in a boiling water bath for 20 min to destroy the enzyme activity then cooled. 100 μl of saline buffer were added to each tube (control and treatment) and absorbance was read at 405 nm with a spectrophotometer (UNICO, SP2100 UV, China) for approximately 5 min, then the absorbency/ min (total activity) was obtained by using the maximum linear rate for both the test and blank.

Specific enzyme activity as unit/min/ml protein was calculated according to the following equation:

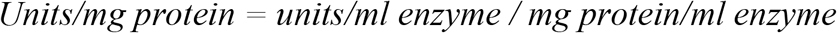

#### Purification of fat body lipase

Purification of the tissue lipase extracted from the fat body was performed in the two steps based on a procedure described by Orscelk et al. [14], with some modifications:

#### Ammonium sulfate [(NH_4_)_2_SO_4_] precipitation

Samples were first subjected to ammonium sulfate precipitation by using 40 and 80% of ammonium sulfate solution and the ammonium sulfate fraction was then collected and centrifuged at 10,000 rpm for 20 min. All the precipitation steps were carried out at 4°C, and in each step, the enzyme activity and protein content were determined.

#### Sephadex G-100 gel filtration chromatography

The last ammonium sulfate fraction was subjected to gel filtration on a Dried Sephadex G-100 column. The dried gel in sufficient amount was incubated in distilled water for 5 h at 90°C. After cooling and removal of air in the gel; it was loaded onto the column (12 × 2 cm) at 27°C. Then the column was equilibrated with 20 mM universal buffer (pH 10), containing 50 mM ammonium sulfate. Enzyme fractions of 3 ml were collected at a flow rate of 20 ml/h with the same buffer. Protein content and lipase activity were measured so that fractions showing the highest activities were pooled for the final step. Sephadex G-100 was obtained from Pharmacia Fine Chemicals, and experiment applied at Regional Center for Mycology and Biotechnology, Al-Azhar University, Cairo, Egypt.

Purification fold and yield% was calculated according to the following equations:

Purification fold = specific activity of purified enzyme/ specific activity of unpurified enzyme.

Yield % = (Total activity of purified enzyme/ total activity of unpurified enzyme)* 100

### Determination of molecular weight and purity of the purified lipases (electrophoretic analysis)

The purity and molecular weight of the purified enzyme was determined by using sodium dodecyl sulfate polyacrylamide gel electrophoresis (SDS-PAGE), as described by Laemmli [23]. A 4% stacking gel and 15% resolving gel were used.

The molecular mass of the enzyme was estimated using BL Uelf Prestained Protein Ladder (from Genedirex) as molecular mass standards (marker proteins). The gel was scanned with gel documentation system by using a scanner and then, the bands were analyzed by using software: Gel-Pro Analyzer ver. 6.0.

### Zymogram analysis

Zymogram analysis of lipase was carried out according to methods of Prim et al. [24], using MUF–butyrate (from Sigma) as the substrate. SDS-PAGE was performed using 15% resolving and 4% stacking gel. By ending of the run the gel gently separated from glasses and immediately rinsed with distilled water then incubated at room temperature in Triton X-100 (2.5% v/v), allowing the renaturation of the enzyme. After 30 min, gels were rinsed again with distilled water and incubated in 100 ml MUF-butyrate solution (100 μM in 50mM phosphate buffer at pH 8.0). After 10 min, gel was put on a UV trans- illuminator to observe fluorescent bands in dark background.

### Determination of biochemical characteristics of the purified lipases

#### 1. Effect of pH on lipase activity

The effect of pH on the purified lipase activity was measured by using lipase diluted in 15 μl universal buffer [25]. Buffers were prepared for the pH range from 2 to 13. After incubation for 1 h at each pH value, lipase activity was assayed as described above.

#### 2. Effect of temperature on lipase activity

Lipase was diluted in 15 μl buffer (50mM Tris-HCl at pH 7 – 7.5) then incubated for 1 h at temperatures ranging from 20 to 70°C. Immediately after incubation, lipase activity was measured by using pNPB as the substrate, as described earlier.

#### 3. Effect of mono- and di-valent cations on lipase activity

The effect of various ions (CaCl_2_, NaCl and KCl) on lipase activity was determined. 50 μl of buffer solution containing one concentration of ions (0, 10, 20, 30 and 40 mM) along with 30 μl of enzyme were pre-incubated for 1 h at pH 7.5 and 37°C. The pre-incubated mixture was added to a solution including 100 μl of universal buffer (pH 7.5). Other steps were carried out as mentioned earlier.

#### 4. Effect of specific inhibitors on lipase activity

The effects of enzyme inhibitors on lipase activity were studied using different concentrations (0, 0.5, 1, 1.5 and 2 mM) of ethylene glycol-bis (β-aminoethylether) N, N, N′, N′-tetraacetic acid (EGTA), Phenylmethylsulfonyl fluoride (PMSF) and Ethylenediamine tetraacetic acid (EDTA). The purified enzyme (30 μl) was incubated with equal volumes of each inhibitor for 10 min at pH 7.5 and 37°C. The mixture was added to a solution including 30 μl of substrate (pNPB, 50 mM). and then the activity was measured as mentioned above.

#### 5. Kinetic parameters measurements

The kinetic parameters (The Michaelis constant, *K*_*m*_ and the maximum velocity *V*_*max*_) were determeined according to Walsh et al. [26]. Final concentrations for *p*-nitrophenol butyrate were 10, 20, 30, 40 and 60 mM, using 30 ul of diluted enzyme preparations at 37°C and pH 7.5 in each assay.

### Statistical analysis

The obtained results were tested using the one-way analysis of variance (ANOVA) followed by Tukey’s test when P ≤ 0.05. Kinetic parameters of the enzyme were estimated by Microsoft Excel.

## Results

### Purification of intracellular fat body lipase from *G*. *mellonella* larvae

The measurements of protein content and lipase activity in crude extract from fat body tissues of *G*. *mellonella* larvae demonstrated that the amount of protein accounted for 1.6702±0.023mg/ml while Spesific lipase activity was 0.016102±0.000278 (μ/mg).

After second ammonium sulfate precipitation, fat body lipase showed a specific activity of 0.0844±0.00216 U/mg protein, 1.68±0.0724 mg/ml of protein, 52.43% of recovery and 5.3 fold purification (Table 1). The ammonium sulfate fractions were then loaded to the Sephadex G-100 column. The sample was fractionated into 50 fractions from which 21 fractions showed lipase activity with three peaks of specific activities at fractions number 6, 8 and 17, and fraction number 6 showed the highest specific activity (Fig 1).

**Table 1:**
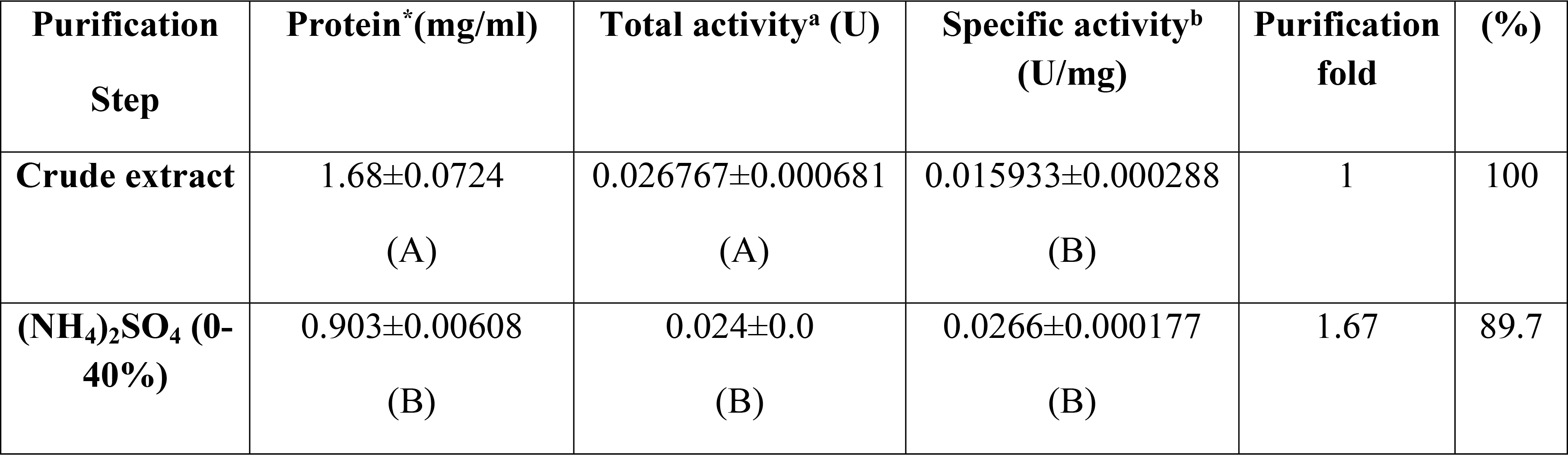
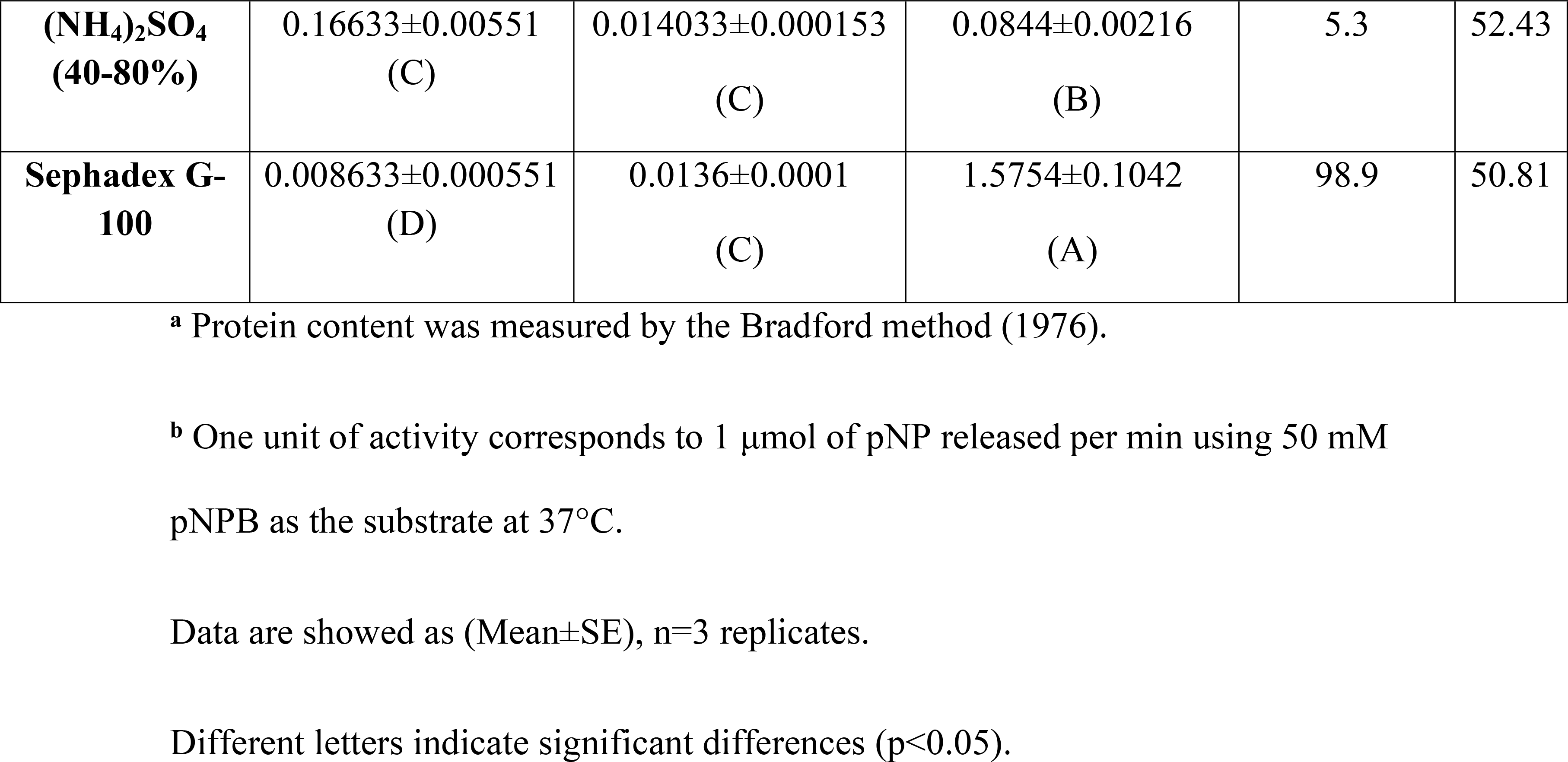
Purification process of fat body lipase from *G*. *mellonella* larvae.

**Fig 1.**
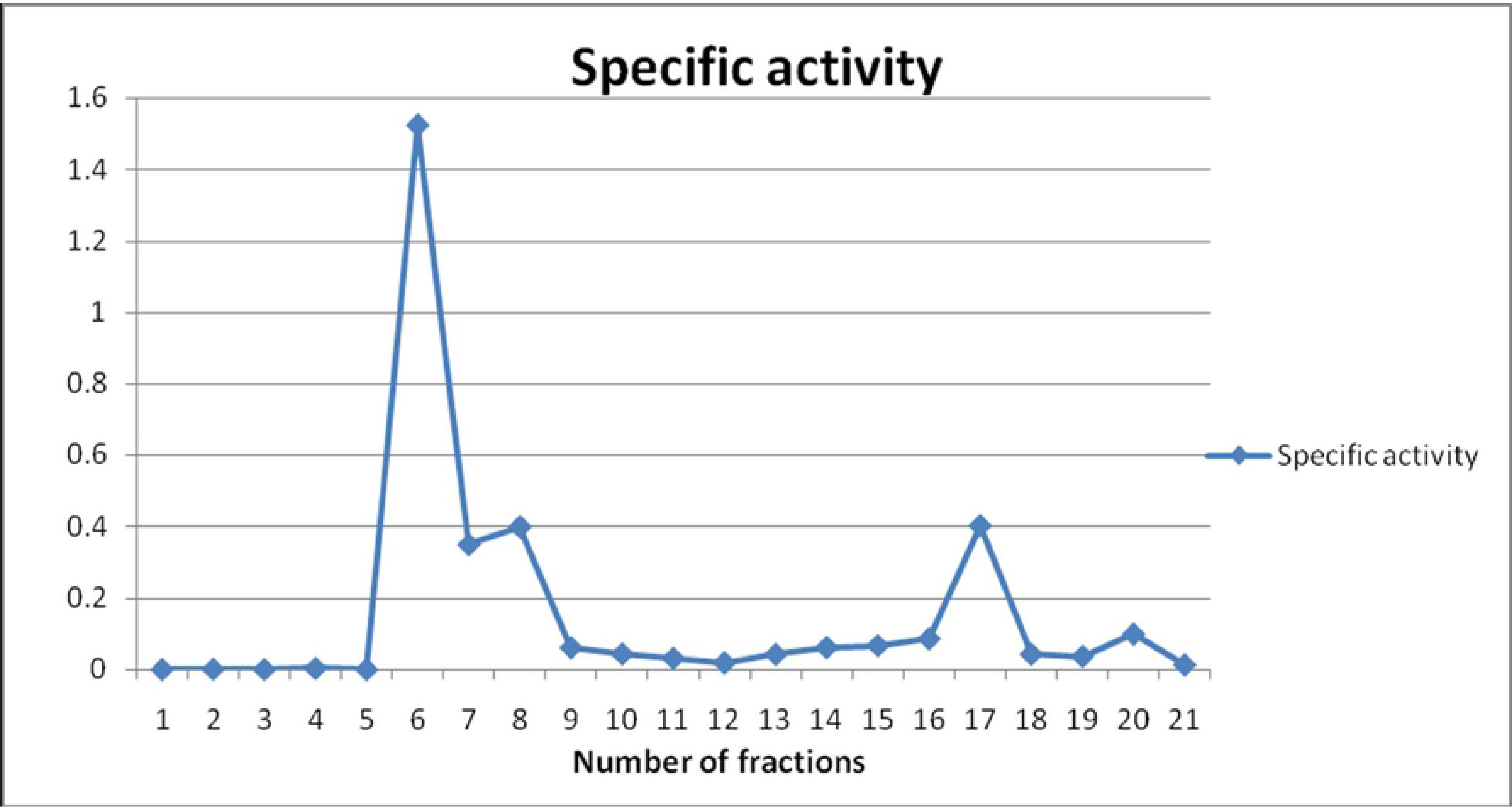
Purification process of fat body lipase from *G*. *mellonella* larvae. Sephadex G −100 gel-filtration of lipase after ammonium sulfate (40 and 80%) treatment. Fractions number 6 to 21 contained the high enzymatic activity on pNPB (50 mM) and fraction number 6 with the highest specific activity (U/mg) was collected for next steps.

After last purification step; the amount of protein and the specific activity of fat body lipase were 0.008633±0.000551 mg/ml and 1.5754±0.1042 μmol/min/mg protein, respectively with 50.81% recovery and 98.9 fold purification. Statistically it was observed that there was no significant difference in specific lipase activity between crude sample and sample after ammunium sulphate percipitation steps but a large significant difference was observed between latter samples and sample that undergo final step of purification (Fig 2). That means the sephadex G-100 step was more effective in purification process.

**Fig 2.**
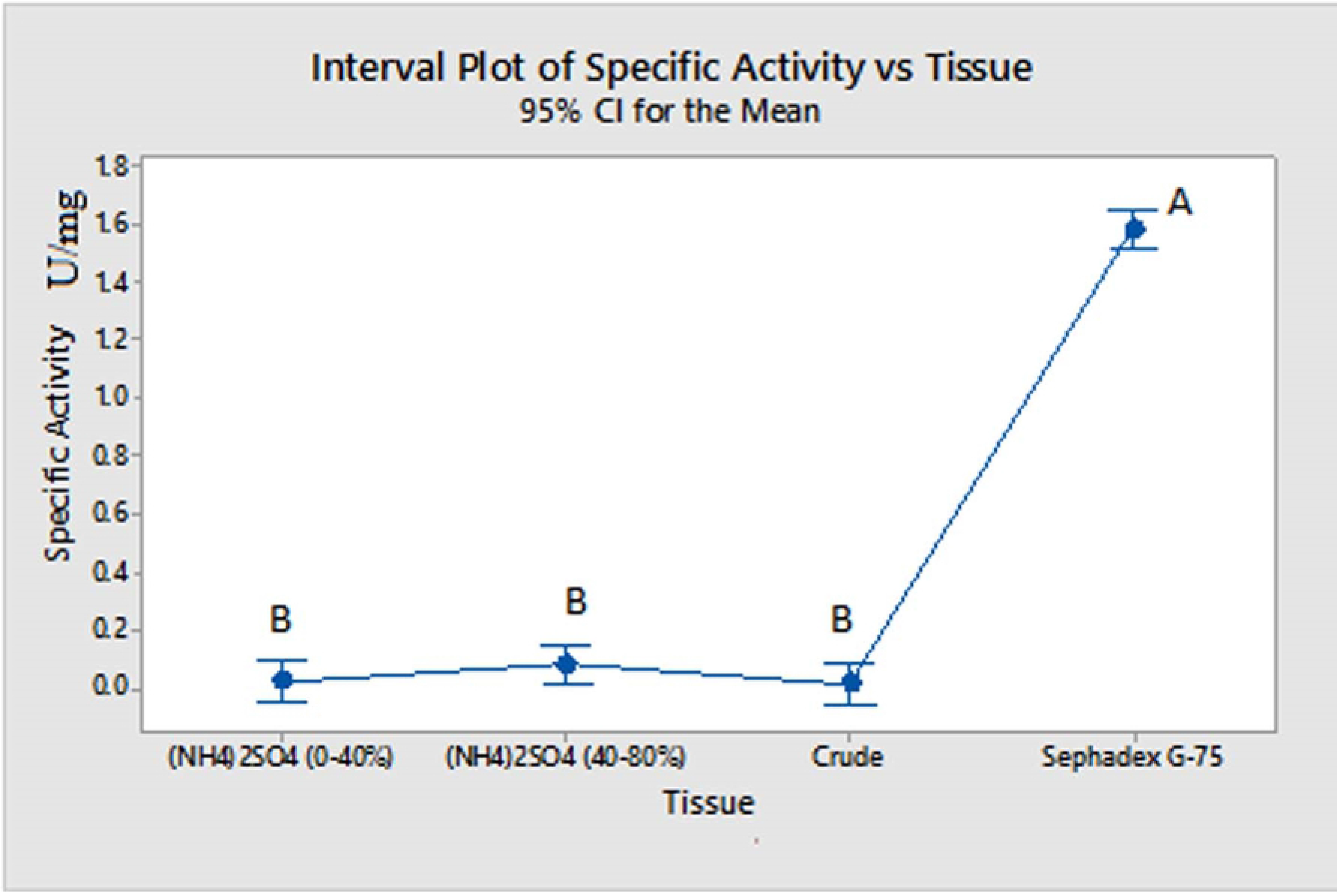
Specific activity of fat body lipase before purification and during steps of purification. Different letters indicate significant differences (p<0.05).

### Determination of molecular weight and purity of the purified lipases from *G*. *mellonella* larvae: SDS-PAGE and Zymogram

SDS-PAGE showed that a number of protein bands were eliminated after each purification step and the lowest number of bands that appeared on gel were after final step (Fig 3). Zymogram analysis yielded two bands with molecular weights of 178.8 kDa and 62.6 kDa for fat body (Fig 4).

**Fig 3.**
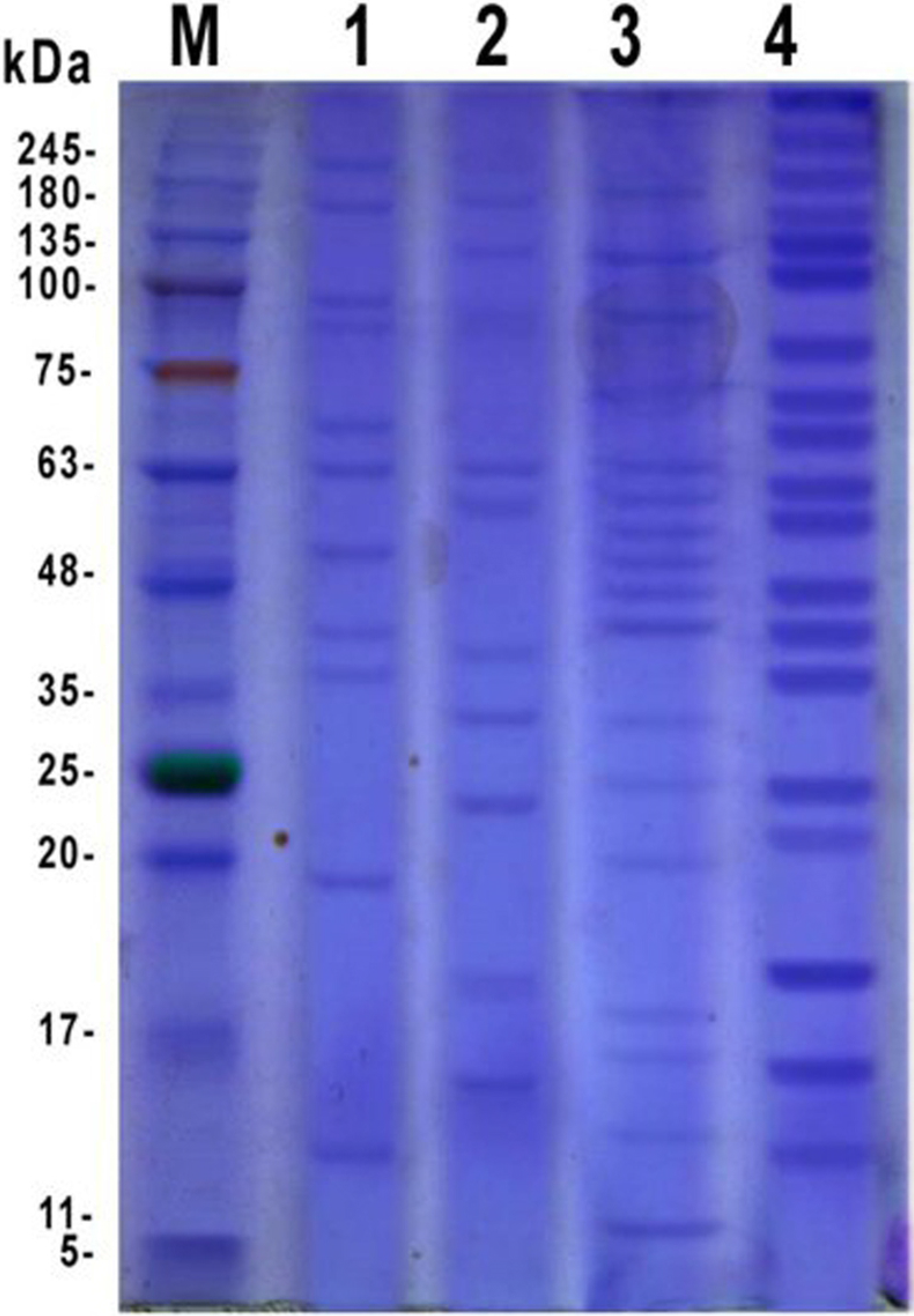
SDS-PAGE of purified fat body lipase from *G*. *mellonella* larvae. M: Molecular marker, 1: Fraction 17 after sephadex G-100 column, 2: Fraction 6 after loading column, 3: Sample after amm. sulfate percipitation (40-80%), 4: Unpurified fat body extract.

**Fig 4.**
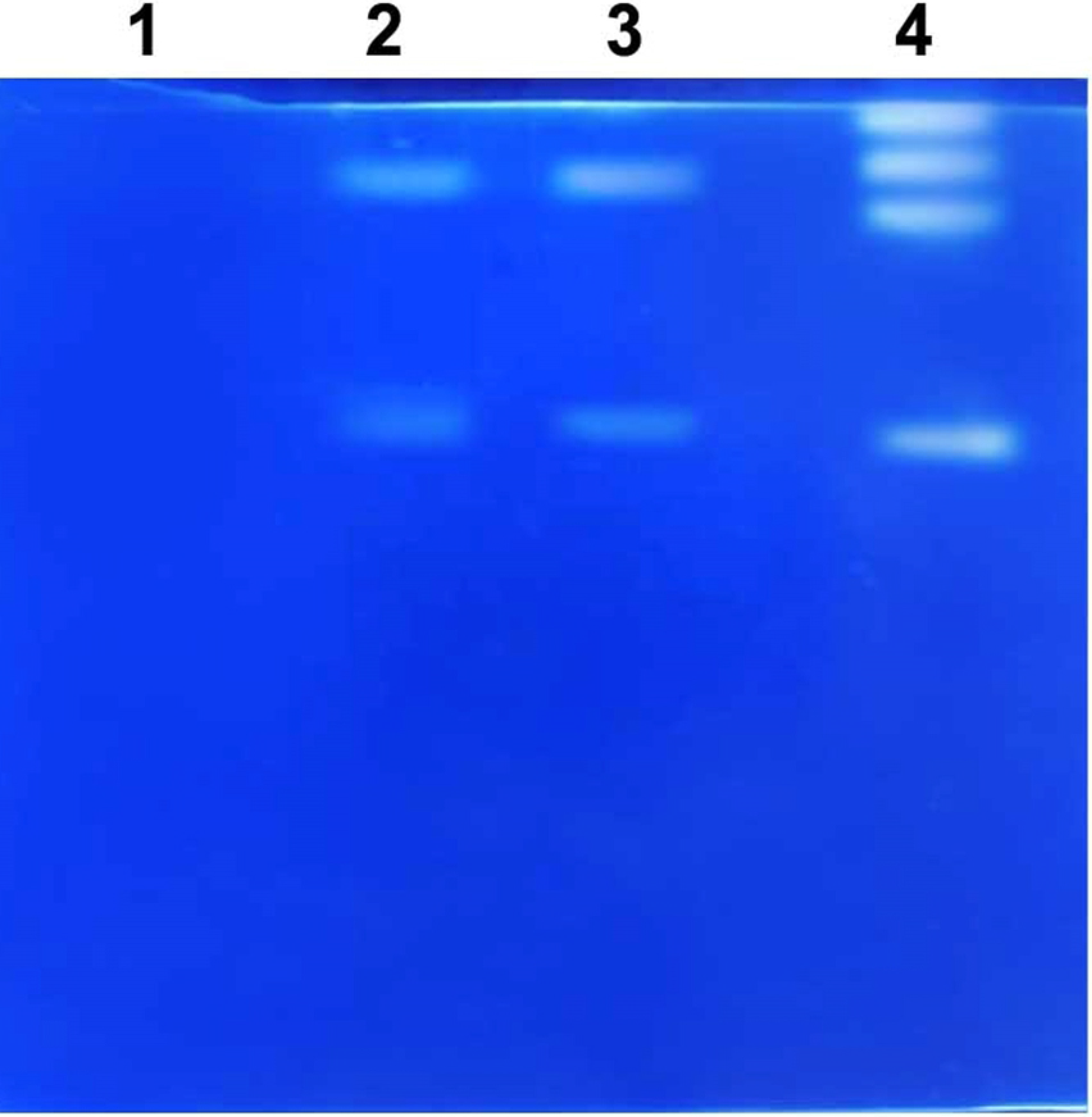
Zymogram of purified fat body lipase from *G*. *mellonella* larvae. 1: Fraction 17 after sephadex G-100 column, 2: Fraction 6 after loading column, 3: Sample after amm. sulfate percipitation (40-80%), 4: Unpurified fat body extract.

### Biochemical characterization of purified intracellular fat body lipase

#### 1. The effect of pH on lipase activity

The effect of pH on lipase activity is presented in (Fig 5) with significant differences (*p*<0.05) were observed. F-values were 379.23. The enzyme activity steadily increased from 4 to 8 and then decreased until pH 13. Activity was high when assayed at pH 7–10, with the maximal activity at pH 8. Lipase activity was decreased at pH 3.

**Fig 5.**
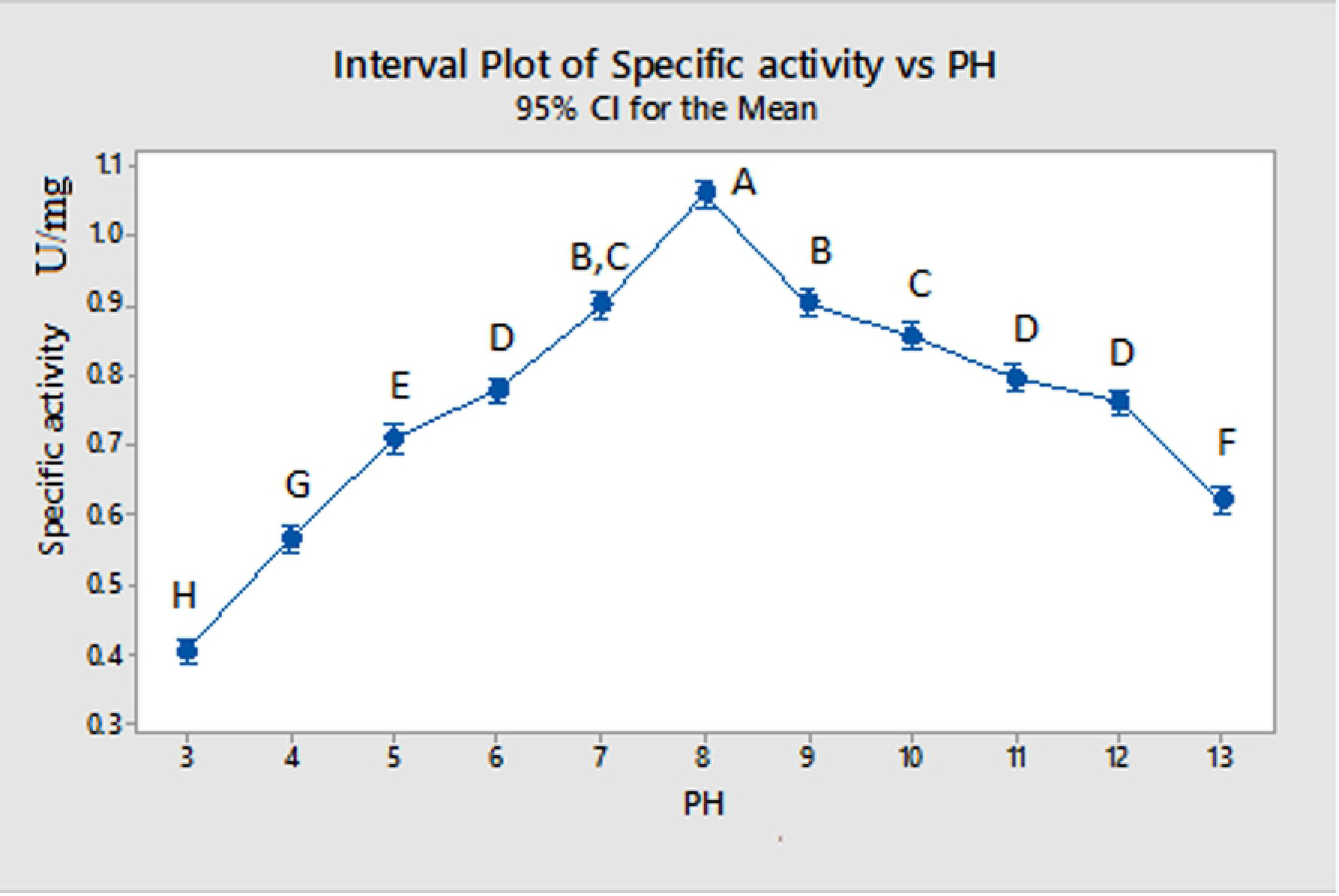
Effect of pH on activity of purified fat body lipase from larvae of *G*. *mellonella* by using different pH range. Different letters indicate significant differences (p<0.05).

#### 2. The effect of temperature on lipase activity

The effect of the temperature on the purified lipase activity was assayed over a range from 20 to 70°C (Fig 6). The purified lipase showed a steady increase in its activity by elevating of the incubation temperature from 20-40°C and decreased till 70°C. Maximum activity under these conditions was at both 37 and 40°C. The statistical analysis demonstrated that highest activity of the enzyme at 37 and 40°C were similar to each other (*p*≤0.05) (F = 60.81).

**Fig 6.**
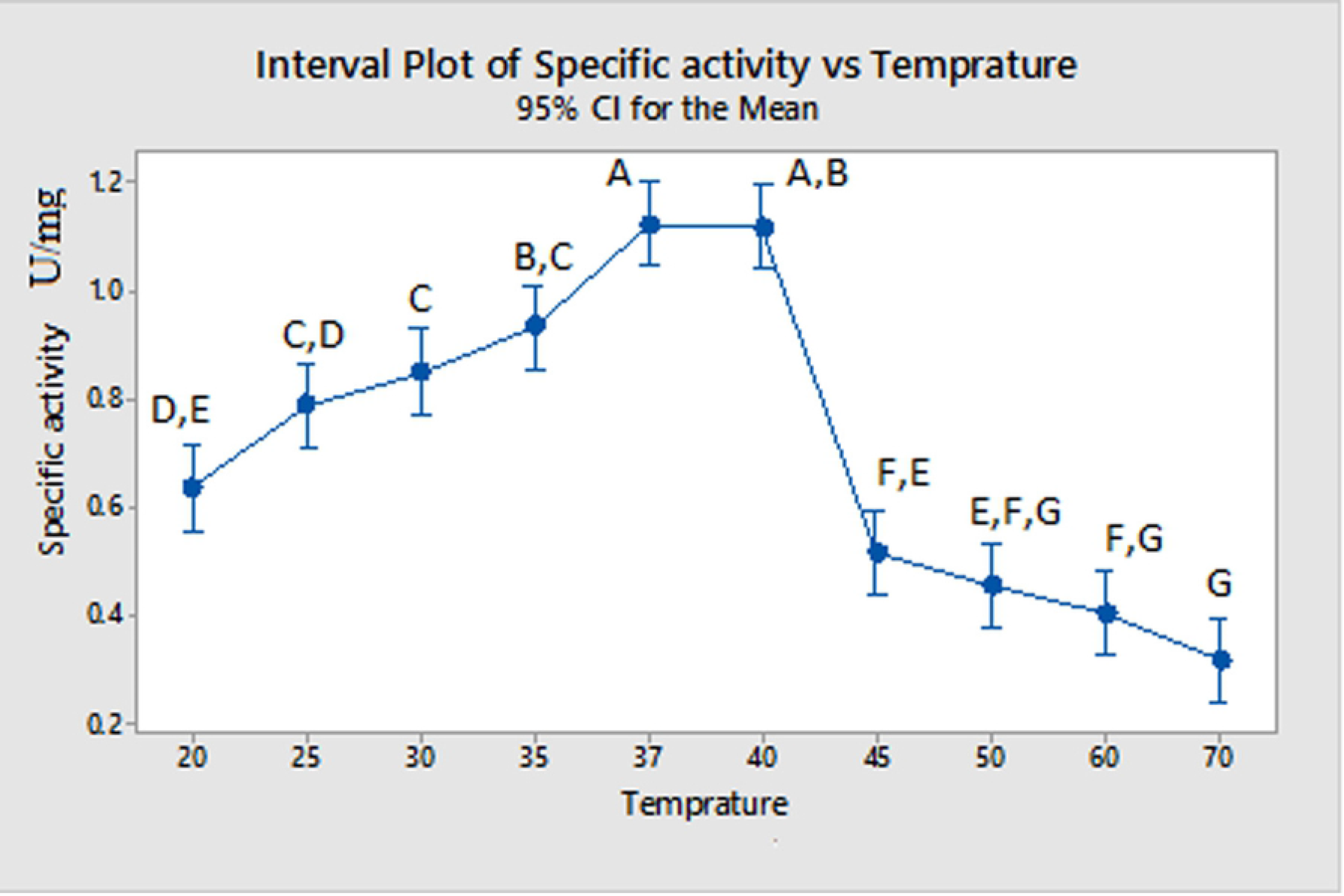
Effect of temperature (°C) on activity of purified fat body lipase from larvae of *G*. *mellonella* by using different temperature range. Different letters indicate significant differences (p<0.05).

#### 3. Effect of mono- and di-valent cations on fat body lipase activity

Effect of mono- and di-valent cations (in several concentrations) on activity of the purified lipase is shown in (Fig 7). All different concentrations of Ca^2+^ significantly (*p*≤0.05) increased the fat body lipase activity so that the activity of enzyme was 2.918±0.0746 μmol/min/mg protein in the highest concentration that is 2.144-fold more than of the enzyme activity without using Ca^2+^. Effect of Na^+^ and K^+^ show significant increase (*p* <0.05) from 0 concentration of Na^+^ and K^+^ to 40 mM concentration. F-values of fat body lipase tests were 104.95, 47.90 and 83.25 for Ca^2^, Na^+^ and K^+^ respectively.

**Fig 7.**
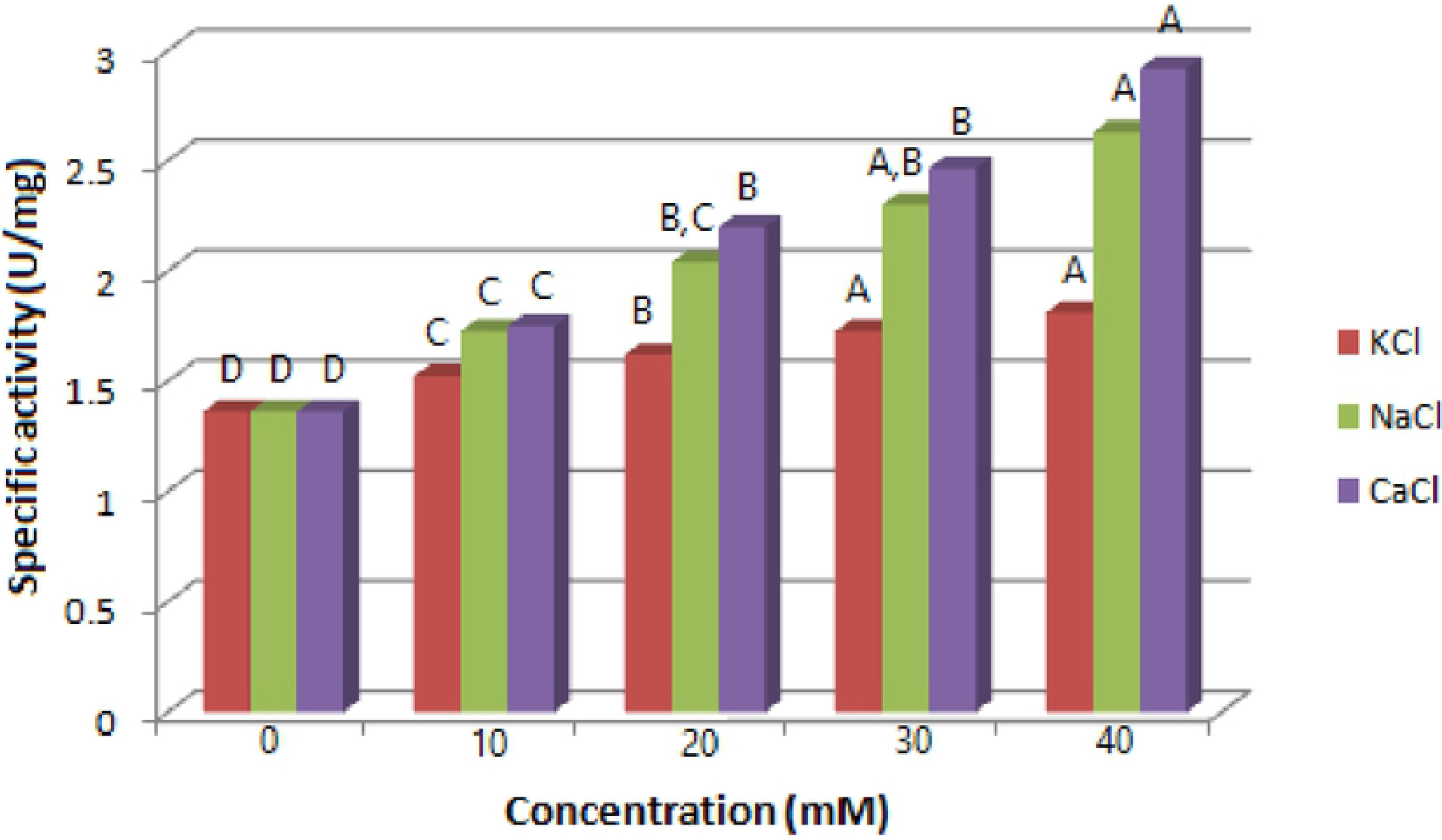
Effect of KCl, NaCl and CaCl_2_ on purified lipase from larval fat body tissue of *G*. *mellonella*. All experiments were carried out at 37°C. pNPB (50 mM) was used as substrate. The enzyme reaction was incubated with ions separately for 1 h. Different letters indicate significant differences (p<0.05).

#### 4. Effect of specific inhibitors on lipase activity of *G*. *mellonella*

PMSF (serine protease inhibitor), EGTA (calcium specific chelating agents) and EDTA (general chelating agent) were used to finding role of metal ions and serine residue in the active site of fat body lipase (Fig 8). It was obtained that different concentrations of PMSF had significant inhibitory effect on lipase from fat body with sharp decrease in activity at 0.5 mM concentration of inhibitor in comparison with control. For EGTA it was found that all tested concentrations decrease enzyme activity. Also EDTA has a significant gradual inhibitory effect on fat body lipase with the lowest activity at concentration of 2 mM. Results showed gradually reduction of purified lipase activity by increasing concentrations of PMSF, EGTA and EDTA indicating the presence of metal ions specially Ca^2+^ and a serine residue in the active sites of the enzymes. Statistically all tests were significantly different with *p*-value < 0.05.

**Fig 8.**
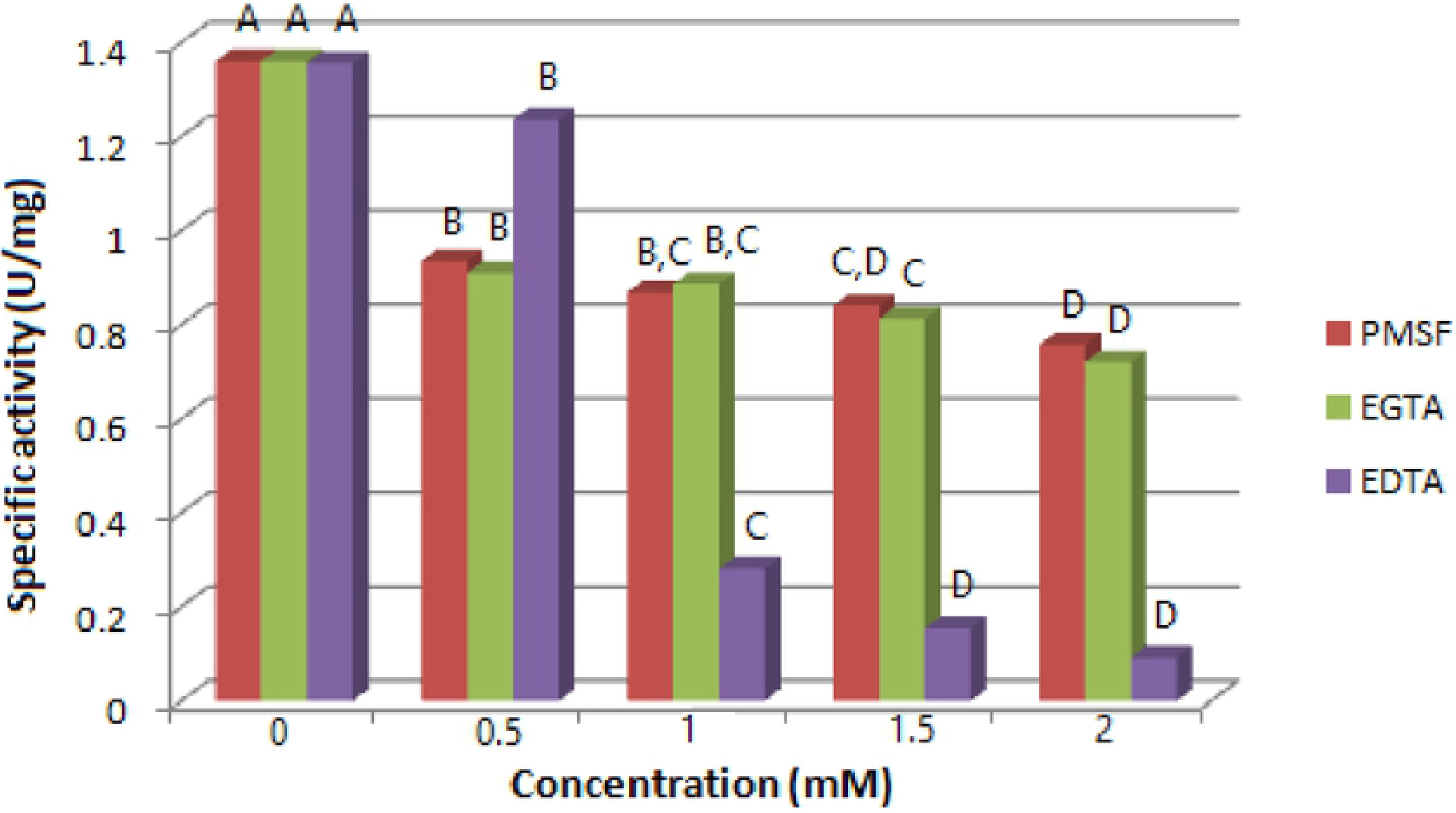
Effect of PMSF, EGTA and EDTA on purified lipase from larval fat body tissue of *G*. *mellonella*. All experiments were carried out at 37°C. pNPB (50 mM) was used as substrate. The enzyme reaction was incubated with various concentrations of inhibitor separately for 10 min. Different letters indicate significant differences (p<0.05).

#### 5. Kinetic parameters of purified lipasee from larval fat body of *G*. *mellonella*

After using several concentrations of p-nitrophenol butyrate to measure kinetic parameters of the purified lipase, the maximum velocity (V_max_) and Michaelis constant (K_m_) were calculated by Microsoft Excel. It was found that; the V_max_ of fat body lipase was 0.316 ± 1.24 U/mg protein while K_m_ was 301.59 ± 24.973 mM (Fig 9). The statistical analysis showed *p*-value of 0.02.

**Fig 9.**
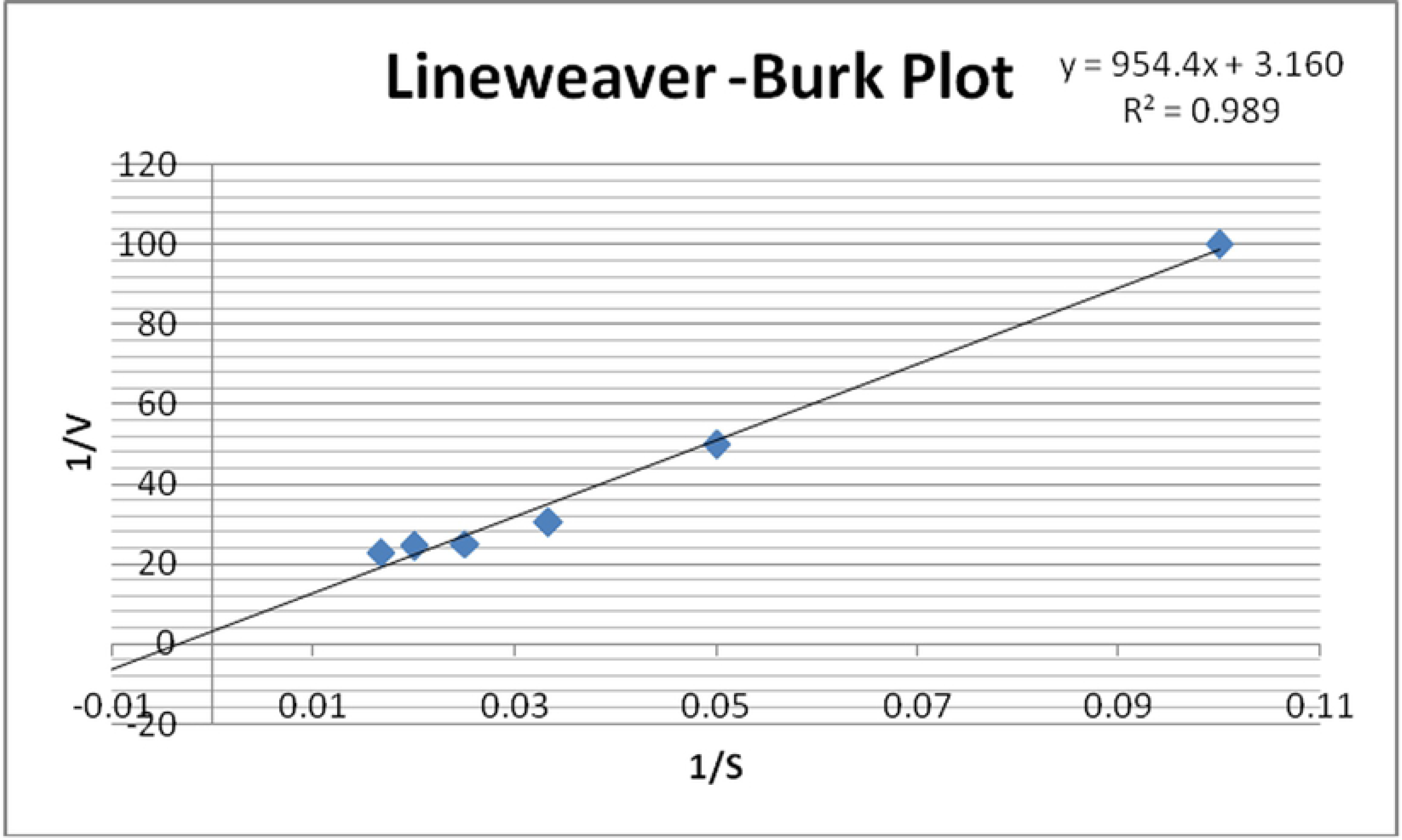
Double reciprocal plot to show the kinetic parameters of purified lipase from the larval fat body tissue of *G*. *mellonella* by using pNPB (from 10 to 60 mM) (1/V_max_= intercept on the 1/V_0_ ordinate, Km/Vmax= the slope of the regression line.

## Discussion

The presented study describes for the first time the possibility to use *G*. *mellonella* larvae for identification of intracellular lipase extracted from the fat body. This study also provides direct tests for characterization of enzyme properties by performing purification and biochemical approaches, which is an advantage for studies related to physiology and pest control.

The extracted lipase from larval and fat body was purified by using two steps. Non- protein components were separated from proteins using ammonium sulfate in a salting out process. Adding ammonium sulfate decreased solvent molecules which interact with proteins by interacting with them [27]. Some proteins coagulate as precipitate because of decreasing the number of solvent molecules. Also separation of a specific enzyme from other proteins can be done through this step.

A gel filtration followed the previous step which separate large proteins according to molecular weights. Very large proteins were excluded while the smaller ones entered between gel particles. Larger protein molecules path down the column and recovered with shorter elution time [27]. It was found that the selected procedures for purification of the larval fat body lipase were effective to eliminate some of contaminating proteins and non-protein molecules so that the final preparation was found to be electrophoretically more partially pure than crude by showing few number bands on SDS-PAGE.

Various purification methods depend on non-specific techniques, precipitation, gel filtration, ion exchange chromatography, affinity chromatography and hydrophobic interaction chromatography have been used to isolate and purify different lipases. Orscelk et al. [14] has purified total body lipase from *Gryllus campestris* by ammonium sulfate precipitation followed by gel filtration. Ranjbar et al. [28] has purified digestive lipases was purified from several insects by three purification steps; such as *Ectomyelois ceratoniae* [28]; *Naranga aenescens by* Zibaee et al. [16] and from *Antheraea drury* by Marepally and Benarjee [29].

Electrophoretic analysis was carried out under reducing conditions for protein profile and zymogram, respectively. For reducing conditions, samples were diluted (1:2) with a sample buffer containing sodium dodecyl sulfate (SDS), glycerol, bromophenol blue and β mercaptoethanol and heated for 2 min at 100°C.

In the current study protein profiles were resolved by (SDS-PAGE). Electrophoretic separation was performed at a 15 mA constant current and at 2°C. Coomassie brilliant blue R-250 was used to visualize the separated proteins. In our study the analyzed fractions show fewer numbers of bands than crude by SDS-PAGE that is due to elimination of unwanted proteins from fractions through purification processing. Invertebrate lipases vary widely in their molecular mass, zymographic analysis showed that intracellular fat body lipase consisted of two monomers with molecular mass of 178.8 kDa and 62.2 kDa. Indeed, nature of enzyme vary with food and organism from which they extracted [30, 31, 32, 33, 34]. Other molecular masses were observer for many insects such 76 kDa for fat body lipase of *M*. *sexta* [6], 30 kDa for lipase of *Cephaloleia presignis* [35], 28 kDa for lipase from digestive juices of *Bombex mori* [36] and 196 kDa for intracellular lipase from pleopods of *Litopenaeus vannamei* [37].

The pH is one of the most important factors which biochemical reactions depend on. Fat body lipase showed the greatest activity at pH 8 followed by 9 while the most activity was lost at pH 3. It seems that lipases from *G*. *mellonella* have a slightly alkaline optimum pH, these results are in conscience with that of kissing bug, *R*. *prolixus* which have high lipase activity at pH 7.0–7.5 [38], and tobacco hornworm *M*. *sexta* at pH 7.9 [6].

The high activity of enzyme at certain pH gives indication on the pH of the environment where the organism feeds [38]. The pH affects the enzyme activity by altering the charge of substrate or of enzyme active site. Very high and low pH can cause electrostatic repulsion [39]. Extremes in pH can also interrupt the hydrogen bonds that hold the enzyme in its three-dimensional structure [32]. However, these changes may be canceled if enzyme returned back to its optimal conditions [27].

Another factor that biochemical reactions rely on is temperature. In the current study, the effect of the temperature on lipase activity was assayed over a range from 20 to 70°C. The maximum activity for fat body intracellular lipase was at 37-40°C while the minimum activity was at 70°C. It was concluded that the enzyme activity reached its peak at the most suitable range of temperature then this activity decreased by increasing the temperature above the suitable one till reached the point at which enzyme denatured then activity decreased sharply. High temperature may interfere with hydrogen bond of enzyme causing denaturation of this enzyme [32]. Similar results have been shown in other insect lipases, as in gypsy moth *Lymantria dispar* [40], *Rhynchophorus palmarum* [41] and *N*. *aenescens* [16]. In contrast, some invertebrates showed highest activity of lipase at 60°C such as the Mediterranean green crab *Carcinus mediterraneus* [42] and the Antarctic krill *Euphasia superba* [43].

In our investigation, it was found that CaCl_2_, NaCl and KCl had an increasing effect on the activity of the partially purified lipase from the fat body of *G. mellonella* larvae. All different concentrations of Ca^2+^, Na^+^ and K^+^ are significantly increased the activity of fat body intracellular lipase.

Similar results were shown for other insects; [38] and Santana et al. [41] showed that the activity of lipase from *R*. *prolixus* and *Rhynchophorus palmarum* increased by increasing calcium ion concentrations respectively. Researches on *C*. *suppressalis* by Zibaee et al. [44] and on *N*. *aenescens* by Zibaee et al. 16] revealed the same conclusion. All previous results indicated that the purified lipase is metalloproteinase, enzyme whose catalytic mechanism involve metal ions, and this result is confirmed by using EDTA and EGTA in current study.

On the other hand, Rivera-Pérez et al. [37] concluded that invertebrate lipases did not depend on a moderate concentration of calcium for maximum activity and stability, as observed in mammalian lipases, e.g., lipase from the adipose tissue showed its maximum activity at 10 mM CaCl_2_. The same conclusion was stated by Mrdakovic et al. [40], on the digestive lipase from gypsy moth, *L. dispar* that is not dependent on Ca^+2^ for activation or stability. These authors depending on the observation of Kim et al. [45] about lipase from *Pseudomona cepacia* which stated that; the stabilization of lipase triad structure is due to the structure of calcium binding site which consists of two carboxylate groups of Asp242 and Asp288 and two carbonyl groups of Gln292 and Val296, taking in consideration that not all invertebrates have this site.

There are several ways by which ions can affect enzyme activity; one of these ways is keeping the enzyme and substrate near to each other to enhance enzyme activity. Also they preclude the unwanted reaction of nucleophiles and put the active groups of enzyme and substrate in the most perfect position leading to increase the ability of enzyme-substrate complex and enzyme stability [41].

Inhibitors are chemicals that diminish enzyme activity. They affect the catalytic characteristics of active site directly or indirectly [27]. In this study three chelating agents were used; EDTA is a general chelating agent, EGTA is a calcium specific chelating agent and PMSF is a serine protease inhibitor. All previous inhibitors were used to finding role of metal ions and serine residue in the active site of enzyme, respectively.

In the current study, the tested synthetic inhibitors had a significant reduction on the lipase activity purified from the fat body of *G*. *mellonella* larvae. The previous results indicated that fat body lipase requires metal ions specially calcium, and have a serine residue at their active site. The effect of PMSF was also shown in TAG-lipase activity from the reproductive accessory glands of the sand fly *Phlebotomus papatasi* [46] and *Rhynchophorus palmarum* [41]. Also it was proven that the active-site serine residue is part of a preserved sequence (GXSXG) that has been found in most lipase sequences of mammals and prokaryotes [47, 48, 49] and invertebrate lipases such as insects [2, 38] and crustaceae [50].

It is important to determine the kinetic parameters of enzyme as it gives us an important information about enzyme behavior and efficiency. The reaction rate is measured and the effects of varying the conditions of the reaction are investigated [26]. Studying enzyme kinetics from this point of view can provide us with information about mechanism by which enzyme catalyze reaction, metabolic role of enzyme and how enzyme activity can be controlled or inhibited using agonist or drugs [51]. K_m_ is the substrate concentration at which enzyme reaches to halfway of its maximal velocity [52]. K_m_ values of enzymes that activate a specific reaction vary according to the organism from which these enzymes were derived.

In the present study, K_m_ values for both fat body lipase was high. High K_m_ values indicate that the enzyme has a low affinity to the substrate. In other word, enzyme will not be saturated with low concentrations of substrate and needs high concentrations to reach its maximal velocity so enzyme activity depends on substrate concentration [27]. At high substrate concentrations; enzyme reaction rate may reaches to its theoretical maximal rate, saturation of enzyme achieved by occupying all of their active sites with substrates and the determination of reaction rate achieved by the intrinsic turnover rate of enzyme [53].

For enzyme that works on several substrates, not in all cases, the lowest K_m_ to a certain substrate supposes that this substrate may be the enzyme’s natural substrate [27]. By taking in consideration that the *G*. *mellonella* was feed on an artificial diet and the nature of secreted enzymes depends on meal on which individual feed on [30, 31, 33, 34, 54], it’s supposed that the p-nitrophenyl butyrate is not the natural substrate of *G*. *mellonella* lipase according to calculated K_m_ value as the low K_m_ value.

## Conclusion

The recent characterization of the intracellular lipase of the wax moth *(G*. *mellonella)* is a major advancement because relatively little was known regarding the enzymes that catalyze the hydrolysis of stored TAGs in insects. The presence of the intracellular lipase in different body tissues in insect larva could be related to energy demands for growth and development. The availability of pure intracellular lipase of larva from the fat body, will permit additional studies on the mechanism of the hormonal regulation of this important metabolic pathway in insects, including an investigation of the role of phosphorylation of the enzyme. This information will provide clues to understanding changes in the lipid catabolism during fasting periods that insects show during molting.

In addition, due to wide hazards resulting from spectrum usage of synthetic chemical pesticides, integrated pest management programs have been focused on the using of enzyme inhibitors in controlling insect pests. The inhibition of lipase could not support the growth and development of *G*. *mellonella* successfully with most unsuitable impact on its reproductive potential. For this reason, synthetic and naturally occurring inhibitors of plant origin are widely regarded as the potential candidates for improved pest control. Before considering inhibitors as a control approach, the enzymatic properties of lipases must be described. The current study could be one of the most important studies on *G. mellonella*. By considering the purification of fat body lipase from larvae and using some inhibitors especially ion chelating agents, it was suggested to develop this study by using lipase inhibitors to reach a successful control of *G*. *mellonella* in the near future.

## Authors Contributions

Rahma R.Z. Mahdy: had performed the practical section of this work, analyzed the data and revised the manuscript.

Shaimaa A. Mo’men: had written and revised the manuscript. Marah M. Abd El-Bar: had written and revised the manuscript.

Emad M.S. Barakat: had designed the work, analyzed the data, written and revised the manuscript

All authors had read the manuscript and approved it for submission.

